# Measuring the strength of selection and pollen limitation on flower colour along an urban rural cline in *Hesperis matronalis*

**DOI:** 10.64898/2026.11.07.764899

**Authors:** Katherine G. Maunder, Maura G. McGregor, John R. Stinchcombe, Megan Bontrager

## Abstract

Closely located urban and rural environments can differ in temperature, water availability, species diversity, and more. As a result, organisms in urban environments may experience different selection than those in rural environments. We investigated whether differences in selection might explain a previously identified urban-rural cline in the flower colour of *Hesperis matronalis* (dame’s rocket). To explore whether plants in urban and rural environments are impacted by possible differences in pollinator availability or behavior, we performed a pollen supplementation experiment. We did not detect an effect of pollen supplementation on seed production in either setting (urban or rural), indicating that there is limited opportunity for any differences in pollinator availability or visitation to explain the cline in flower colour. We also measured the strength of selection acting on flower colour to investigate if it differed between urban and rural populations. In all but one population, we failed to detect significant linear or non-linear selection on flower colour, suggesting that selection on flower colour does not differ between urban and rural populations. Our results suggest that the urban-rural flower colour cline is likely not caused by differences in selection or pollen-limitation and may be due to stochastic forces.

## Introduction

Environments shape selection, and as a result when human activity alters an environment, it can alter selection as well (Des Roches *et al*. 2021). While human activity is changing the environment at a large scale by increasing the global temperature (Al-Ghussain 2019) and the frequency of extreme weather events (Capua and Rahmstorf 2023), it is also changing the environment at more local scales. For instance, urban areas, where human activity is higher, often differ from rural areas in temperature, water permeability, species diversity, habitat connectivity, and resource availability (Kalnay and Cai 2003; Grimm *et al*. 2008; Li *et al*. 2010). As a result, two geographically close populations can face different selective pressures if one is in an urban area while the other is in a rural one. Here we measure the strength of selection on flower petal colour and assess whether pollinators are an important selective pressure in urban and rural populations of an introduced plant species.

Only a limited number of studies have quantitatively measured whether the strength of selection on a given trait differs between urban and rural environments (Charmantier *et al*. 2024). Such quantitative studies have shown that the different selective environments of urban and rural populations can drive variation in flower size and display size (Bode and Tong 2018; Irwin *et al*. 2018). However, no studies have yet quantitatively investigated if differences in urban-rural selective environments might drive variation in flower colour.

Flower colour is a visual trait that can affect how pollinators perceive flowers and the compounds responsible for this colouration also play important roles in stress responses. Flower colour variation is often under selection by pollinators, but it can also be influenced by other drivers (Strauss and Whittall 2007; Caruso *et al*. 2018; Tenhumberg *et al*. 2023), including other biotic factors such as herbivores (Irwin *et al*. 2003; De Jager and Ellis 2014), abiotic factors such as precipitation (Schemske and Bierzychudek 2001; Arista *et al*. 2013), and non-selective processes (Wang *et al*. 2016). Many of these possible drivers of flower colour variation overlap with conditions known to differ between urban and rural populations (e.g. herbivory (Miles *et al*. 2019), moisture (Hao *et al*. 2023), temperature (Phelan *et al*. 2015), etc.). An increasing number of studies have highlighted how urban environments can affect the composition of pollinator communities and plant-pollinator interactions (Harrison and Winfree 2015; Wenzel *et al*. 2020). For instance, in Central Europe the overall diversity of pollinating insect species is lower in urban environments, but the diversity of Hymenoptera species is higher (Theodorou *et al*. 2020). Pollinator behaviour may also differ between urban and rural environments, for example, pollinators are less likely to enter *Robinia pseudoacacia* flowers in urban populations than in rural populations (Buchholz and Kowarik 2019). Changes in pollinator diversity and activity between urban and rural environments may affect selection on flower colour and could drive patterns of phenotypic variation.

An urban-rural cline in flower colour has been previously observed in *Hesperis matronalis* in eastern Ontario, where populations in urban areas have a more diverse mix of flower colour than populations in more rural areas (Maunder *et al*. 2026). Specifically, while urban populations have a relatively even mix of white, pink, and purple flowers, populations in more rural areas consist of mostly purple flowers. In North America, *H. matronalis* is non-native and is the result of garden escape (Francis *et al*. 2009). It is likely that historical processes and non-selective effects during the introductions and subsequent expansions of this species into natural habitats has contributed to current patterns of flower colour variation (Keller and Taylor 2008; McLean *et al*. 2017; Maunder *et al*. 2026). However, the extent to which spatially variable selection may also contribute to this urban-rural cline has not been formally investigated.

We explored how selection on flower petal colour may differ between urban and rural populations of *H. matronalis* in eastern Ontario and investigate the possible contribution of pollinators. If a floral trait affects pollinator preference, trait variation can lead to differences in fitness (Glover and Martin 1998). For example, if white flowers are less attractive than purple flowers to local pollinators, they may be visited less, receive less pollen, and potentially produce fewer seeds. Pollen supplementation can reduce the influence of pollen limitation on seed set and allow researchers to evaluate the extent to which pollinators are potentially mediating selection (Knight *et al*. 2006; Fishman and Willis 2008; Sandring and Ågren 2009). We use a pollen supplementation experiment to ask whether fruits are pollen limited, whether flower petal colour influences pollen limitation, and whether this relationship is impacted by how urban the environment is. Additionally, we quantify the strength of selection on flower petal colour to test whether it differs between more urban and more rural populations. To achieve this, we quantitatively measure flower petal colour, estimate seed production, and calculate both directional and non-directional selection differentials in urban and rural populations (Lande and Arnold 1983).

## Materials and Methods

### Study System

*Hesperis matronalis* is a biennial or short-lived perennial native to Eurasia that was introduced into North America in the 1800s as an ornamental plant (Francis *et al*. 2009). It has become naturalised across the United States and southern Canada and is commonly found along roads, riverbanks, and forest edges (Francis *et al*. 2009)*. Hesperis matronalis* produces flowers that can be either a shade of white, pink, or purple, and while flower colour is consistent within an individual plant, populations of *H. matronalis* are often polymorphic (Maunder *et al*. 2026). Many insects pollinate *H. matronalis*, including bees, butterflies, moths and syrphid flies. Observations of an *H. matronalis* population in Pennsylvania found that syrphid flies made up the majority of pollinators and that visitation was independent of colour (Majetic 2008). It is unclear to what extent insect pollination is required for seed set due to mixed reports of whether *H. matronalis* is self-compatible (Majetic 2008) and possibly predominantly self-fertilising (Susko and Clubb 2008) or obligately outcrossing (Mitchell and Ankeny 2001; Weeks and Frey 2007).

### Site Selection

We sampled *H. matronalis* individuals from ten populations across a gradient of human activity in and around Kingston, Ontario where a cline in flower colour was observed in 2021, 2022, and 2023 (Maunder *et al*. 2026). To identify suitable populations, we screened known naturalised populations of *H. matronalis* from the previous study for sufficient population size (more than 500 plants) and presence of white, pink, and purple colour morphs (target was at least 50 individuals of each morph). To estimate the amount of human activity around populations, we used night sky brightness (NSB) as a proxy (see Maunder *et al*. 2026 for a description of this method). To calculate NSB, we obtained zenith sky brightness values from Falchi *et al*.’s (2016) world atlas of artificial sky luminance (at a spatial resolution of 1 km) and then multiplied these values by negative one. We multiply by negative one to make NSB positively correlated with human activity which is easier to interpret. The night sky brightness of our five more urban populations ranged from -20.79 to -19.17 mag/arcsec^2^, and the night sky brightness of our five more rural populations ranged from -21.89 to -21.37 mag/arcsec^2^.

### Plant Sampling and Field Measurements

Between May 27th and June 7th, 2023, we tagged 250 plants at each site, and measured flower petal colour with a spectrometer. To sample plants, we walked a transect across the site and randomly selected plants every meter by haphazardly pointing a rod without looking. At each site, the same team of researchers tagged selected plants and recorded a visual score of petal colour (white, pink, or purple). We sampled at least 50 plants of each petal colour per population. In populations where any one colour morph was less frequent (where random sampling resulted in less than 50 plants being sampled), off-transect plants of that petal colour were selected to meet the target sample size.

We collected one flower from every sampled plant to quantitatively measure colour. To account for any colour variation due to flower age, we collected all flowers at the same phenological stage, when the anthers were just beginning to emerge from between the petals. After collection, flowers were stored at 4°C for up to 2 hours until they were measured. For every flower, we took three measurements of petal light reflectance, each from a separate petal. We used an SR4 compact spectrometer (Ocean Optics) and a DH-2000 deuterium-tungsten halogen light source (Ocean Optics) to measure light reflectance from 300-700 nm. We did not analyse reflectance at smaller wavelengths because there is no discernible pattern of UV reflection in this species (Maunder *et al*. 2026).

### Pollen Supplementation Experiment

Between June 1^st^ and 8^th,^ we performed a pollen supplementation experiment on plants from six of the populations in the study region (see map in Figure 1B). The pollen supplementation experiment was performed on a subset of flowers instead of the entire plant. At each site, an average of 76 plants (min: 71, max, 80) from the main study were included.

**Figure 1:**
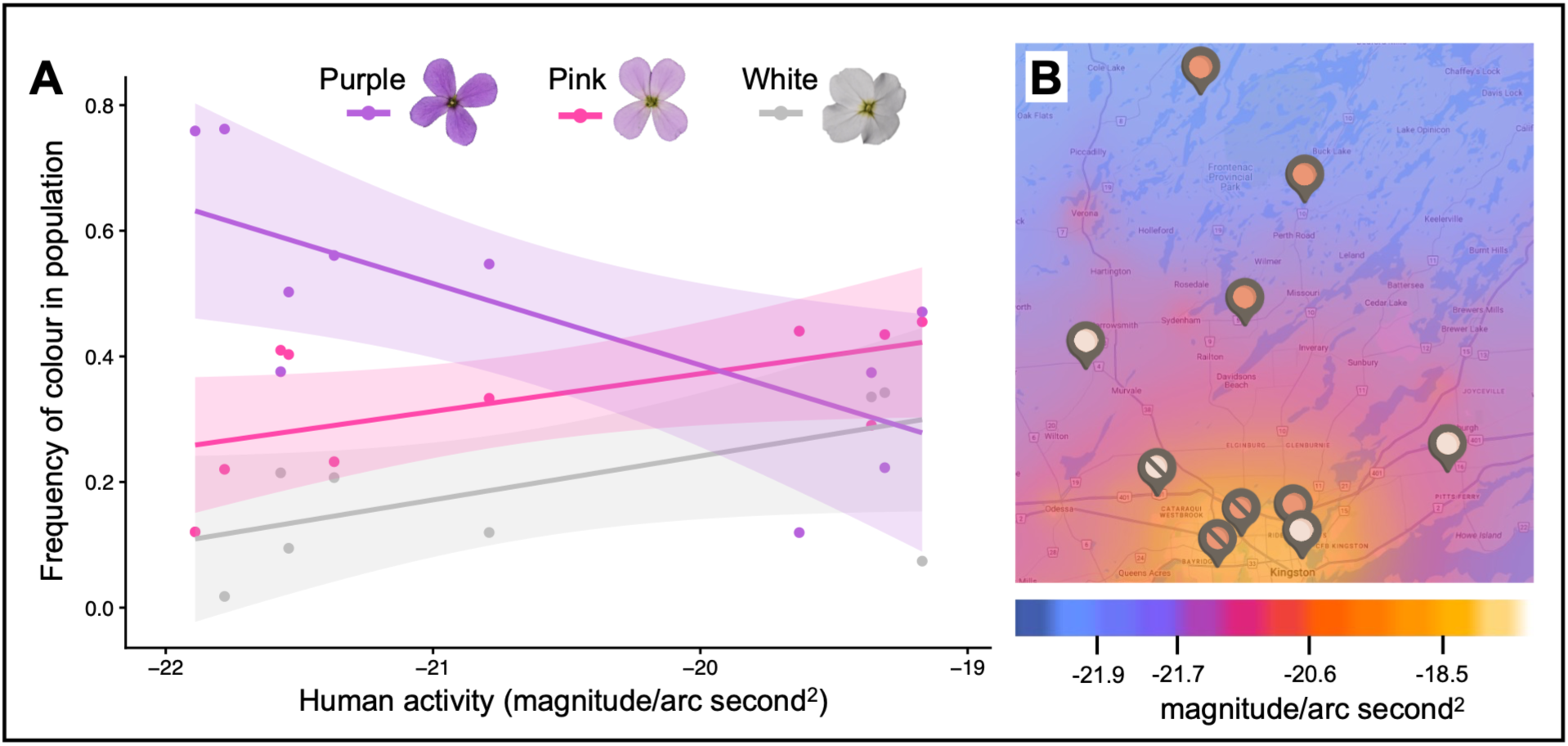
A) Association between the frequency of each flower colour in each population and the amount of human activity in the area with a 95% confidence band. The three flower colours represent visual classifications assigned in the field. B) Population locations on a map of night sky brightness values (magnitude/arc second^2^). We use night sky brightness as an estimate of human activity levels, where a more positive value of night sky brightness represents a higher amount of human activity. Pins indicate the location of each sampled population. Those with orange circles were included in the pollen supplementation experiment, while those with white circles were not. Pins with a slash through the circle identify the three populations where stem number was not recorded.

Approximately half of these plants were given supplemental pollen and half were used as controls, with an average of 13 plants (min 9, max 18) from each visual flower colour category (white, pink, and purple) in each treatment. Pairs of plants with the same flower colour were selected at roughly evenly spaced intervals across the transect and haphazardly assigned to supplementation vs. control treatments. On all plants in both treatments, three flowers in anthesis were selected and marked by placing thin strips of red tape on the pedicels. For plants in the pollen supplementation treatment, we used a paintbrush to pick up pollen from nearby untagged plants of the same flower colour and deposit it on the stigmas of marked flowers. Our pollen supplementation treatment should not affect normal pollinator visitation and should just increase the overall amount of pollen deposited on each stigma. Plants in the control treatment did not receive supplemental pollen.

### Seed Collection and Estimation of Fitness

In mid-August, we collected fruits from tagged plants to measure their reproductive output through seed production. Of the 250 plants tagged in each population 123-223 plants were recovered (mean = 186, median = 191). We aimed to collect fruits late enough in the season that the majority of fruits were developed, but before the fruits opened and dropped their seeds. *Hesperis matronalis* individuals can have one or multiple stems that branch off within 3 cm of the root, at or below the soil level (sampled plants ranged from 1-20 stems, averaging 3). From all tagged plants, one stem was randomly collected, and the total number of stems was recorded (this measurement was not recorded in the first three sites visited due to previous reports of the rarity of multiple stems (Francis *et al*. 2009)). In addition, for plants in the supplemental pollen experiment, marked fruits were harvested and stored separately in coin envelopes. All plant materials were air-dried in paper bags at room temperature for four months. Once dry, we took measurements of plant size and reproductive fitness. For all tagged plants, we measured stem diameter as an estimate of plant size, and for an estimate of fitness, we measured fruit number and the length of 10 randomly selected fruits. We then calculated an estimate of seed set (absolute fitness) by multiplying fruit number by the number of stems on a plant by 2.06 (an estimate for the number of seeds per cm of fruit, see Appendix B Figure 1). For plants in the pollen supplementation treatment, we also measured the length of the three marked fruits, opened these fruits, counted the number of seed indentations along the septum (evidence of seed development (Mitchell and Ankeny 2001)), and counted the number of intact seeds. For all plants in this study, our estimates of fitness represent a potential upper bound of the number of seeds that a plant may have set. By the time of collection, most of the seeds within a fruit were not intact and would easily crumble under light pressure (see Appendix A for an analysis and discussion of seed damage).

### Statistical Analysis

#### Spectral Colour Data and Flower Colour Cline

We used the pavo package (Maia *et al*. 2019) in the R Statistical Software (v4.4.1, R Core Team) to perform locally weighted scatterplot smoothing, using a span of 0.2, on each of the three reflectance spectrums obtained per flower. Principal component analysis revealed that all wavelengths between 430 and 630 nm had a loading weight close to one suggesting that they are highly correlated and that variation in petal colour is largely due to differences in lightness (the amount of light reflected), rather than differences in hue (the specific wavelengths being reflected). Therefore, we used the average reflectance of light between 430 and 630 nm as a simplified univariate measure of colour (moving forward we refer to this value as reflectance). For subsequent analyses we used the median reflectance value (of the three petals measured) to characterize the petal colour of each plant. We ran an ANOVA to compare the reflectance values of the field-assigned flower colour categories (purple, pink, white). To confirm that our sampled populations followed the previously observed urban-rural cline in flower colour frequency we also ran a linear mixed model with reflectance as the response variable, night sky brightness (a proxy of the human activity around each population) as a fixed effect, and population as a random effect.

#### Pollen Supplementation

To examine the relationship between the supplemental pollen treatment and fruit set (whether or not a fruit developed), we used a binomial mixed effect regression with fruit development as the response variable (coded as 0 or 1, with 1 being a developed fruit), pollen treatment (control vs supplemented), reflectance, and their interactions as fixed effects, and plant identity as a random effect (this accounts for the fact that each plant contributed three fruits to these data). Population was not included in the model because some populations had no or very few undeveloped fruits (across all populations only 7% of fruits were undeveloped). To test for an effect of pollen supplementation on the number of seeds produced by individual fruits, we ran a linear mixed effect model with fruit length as the response variable; pollen treatment, population, reflectance, and their interactions as fixed effects; and plant identity as a random effect as before. We used a type III ANOVA using the lmerTest Package (Kuznetsova *et al*. 2017) to investigate the significance of these main and interaction effects.

#### Estimating the Strength of Selection on Flower Colour

To assess if there was a significant relationship between fitness and flower colour, we performed linear and quadratic regressions for each population with relative fitness as the dependent variable and z-score normalised values of reflectance as the explanatory variable (linear: relative fitness ∼ normalized reflectance, quadratic: relative fitness ∼ normalized reflectance + normalized reflectance^2^). Reflectance is the median reflectance value of the three petal measurements. The coefficient of the linear regression represents the total strength of directional selection acting on flower colour (s), and the coefficient of the squared term in the quadratic regression, once multiplied by two, represents the strength of non-linear selection acting on flower colour (C) (Lande and Arnold 1983; Stinchcombe *et al*. 2008).

## Results

### Flower Colour Cline

The flower colour categories assigned in the field (white, pink, and purple) separated plants into groups with significantly different reflectance values (p < 0.01 for all comparisons, Figure 2), though there was some overlap in reflectance values among groups. In line with the previously observed cline (Maunder *et al*. 2026) , we found that the mean reflectance of populations significantly increased as human activity increased (slope = 3.34, SE = 1.31, p < 0.05) and that in areas with low human activity there was an increase in the frequency of plants with low reflectance (purple-coloured flowers) and a decrease in the frequency of plants with mid and high reflectance (pink- and white-coloured flowers) (Figure 1A).

**Figure 2:**
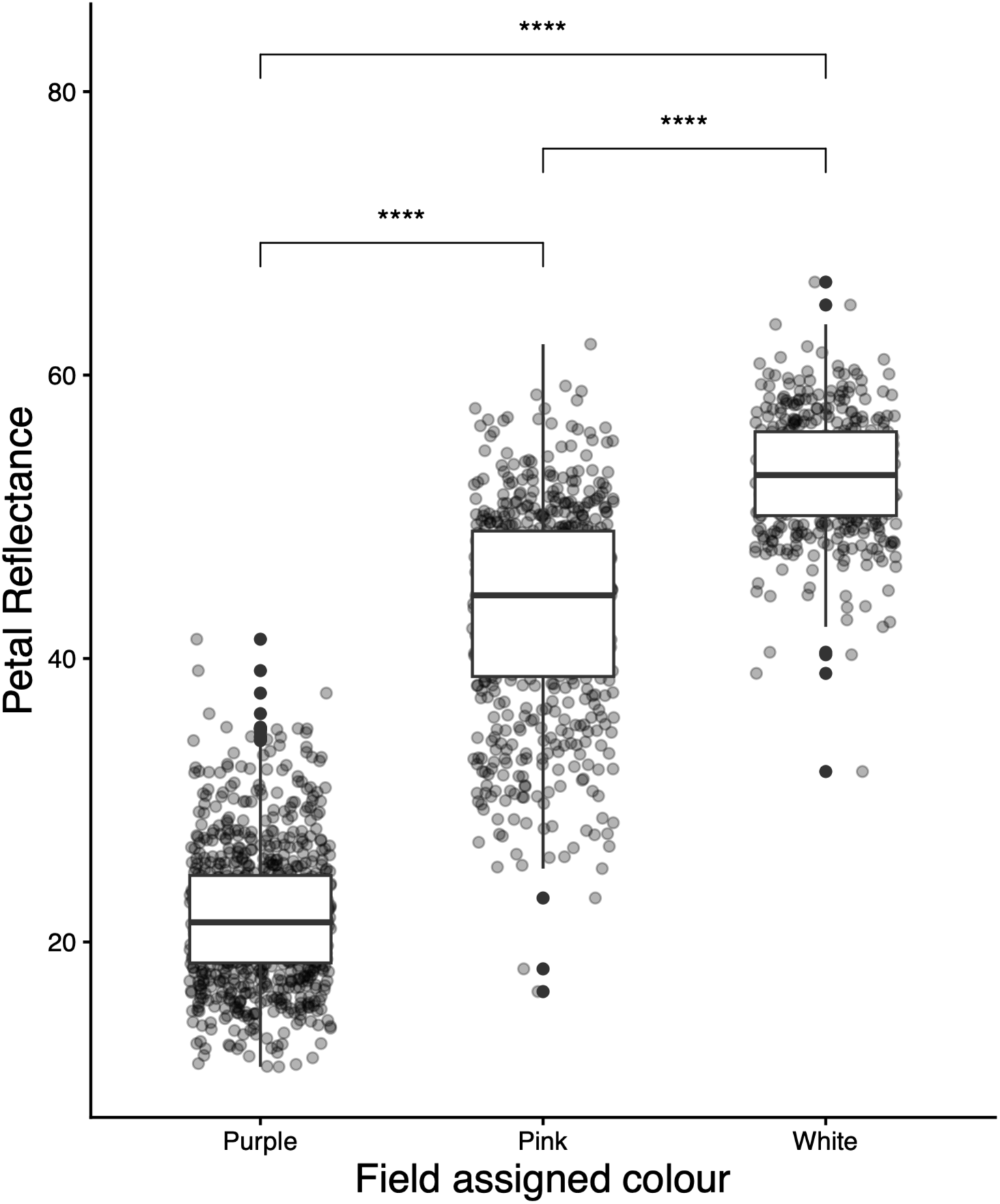
The visual categories of flower colour assigned in the field differ significantly in petal reflectance as measured with the spectrometer (brackets and asterisks indicate significant comparisons). Each point represents the median spectral reflectance measurement of the three petals measured per plant with a horizontal jitter. Petal reflectance refers to the average percentage of light reflected between 430 and 630 nm. Higher reflectance is consistent with paler colouration.

### Pollen Supplementation

We found no significant difference in the probability of fruit set between the pollen supplementation and control treatments (p = 0.439) and no significant effect of the interaction between pollen supplementation treatment and reflectance on fruit set (p = 0.764). In the control treatment 11% of collected fruits were undeveloped (43/395) and in the pollen supplemented treatment 4% of fruits were undeveloped (20/496). Of the fruits that did develop, fruit length was not significantly affected by pollen supplementation (p = 0.843) or the interaction of pollen supplementation with either population (p = 0.793) or reflectance (p = 0.442) (Figure 3). These results collectively suggest that there is no evidence of pollen limitation on fruit set or fruit length.

**Figure 3:**
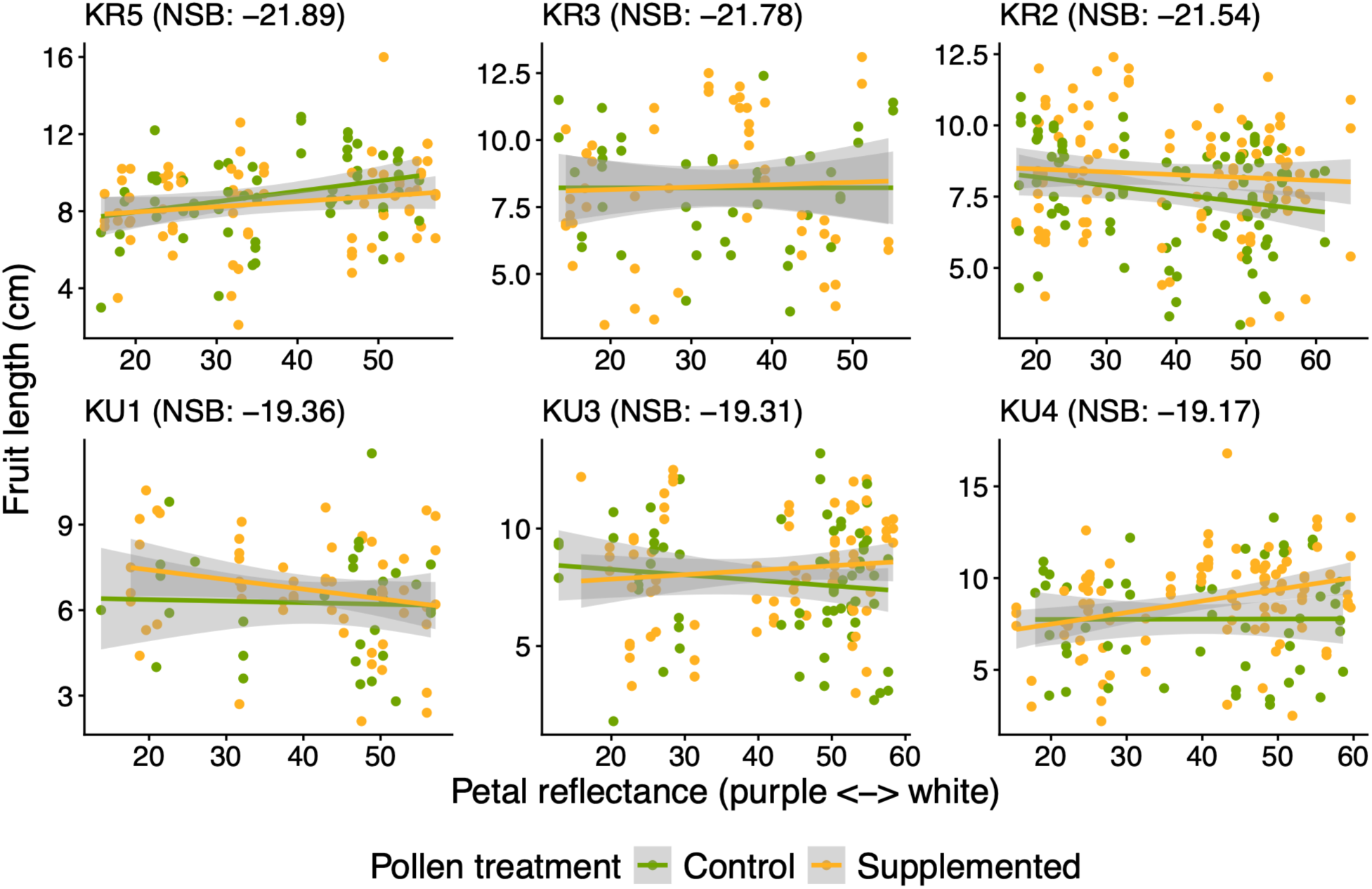
No significant difference between control plants and those that received supplemental pollen in the relationship between fruit length and petal reflectance. Linear regression lines are shown with 95% confidence bands for each treatment (control = green and pollen supplemented = yellow) regardless of whether the regression is significant. An individual plant may be represented in the plot by as many as three points (the three flowers/fruit that were included in the experiment). Night sky brightness (NSB) is an estimate of human activity levels, where a more positive value represents a greater amount of human activity.

### Strength of Selection on Flower Colour

We did not find a significant linear relationship between flower colour and fitness in any population, meaning that we did not find evidence of directional selection (Table 1, Figure 4). There was only a significant quadratic relationship between flower colour and fitness in one of the seven population, indicating a lack of evidence for disruptive or stabilizing selection in most populations (Table 1). The one population with a significant relationship was KR4 where intermediate flower reflectance was associated with lower relative fitness, providing some evidence for disruptive selection (C = 0.66, SE = 0.22, p = 0.0036).

**Figure 4:**
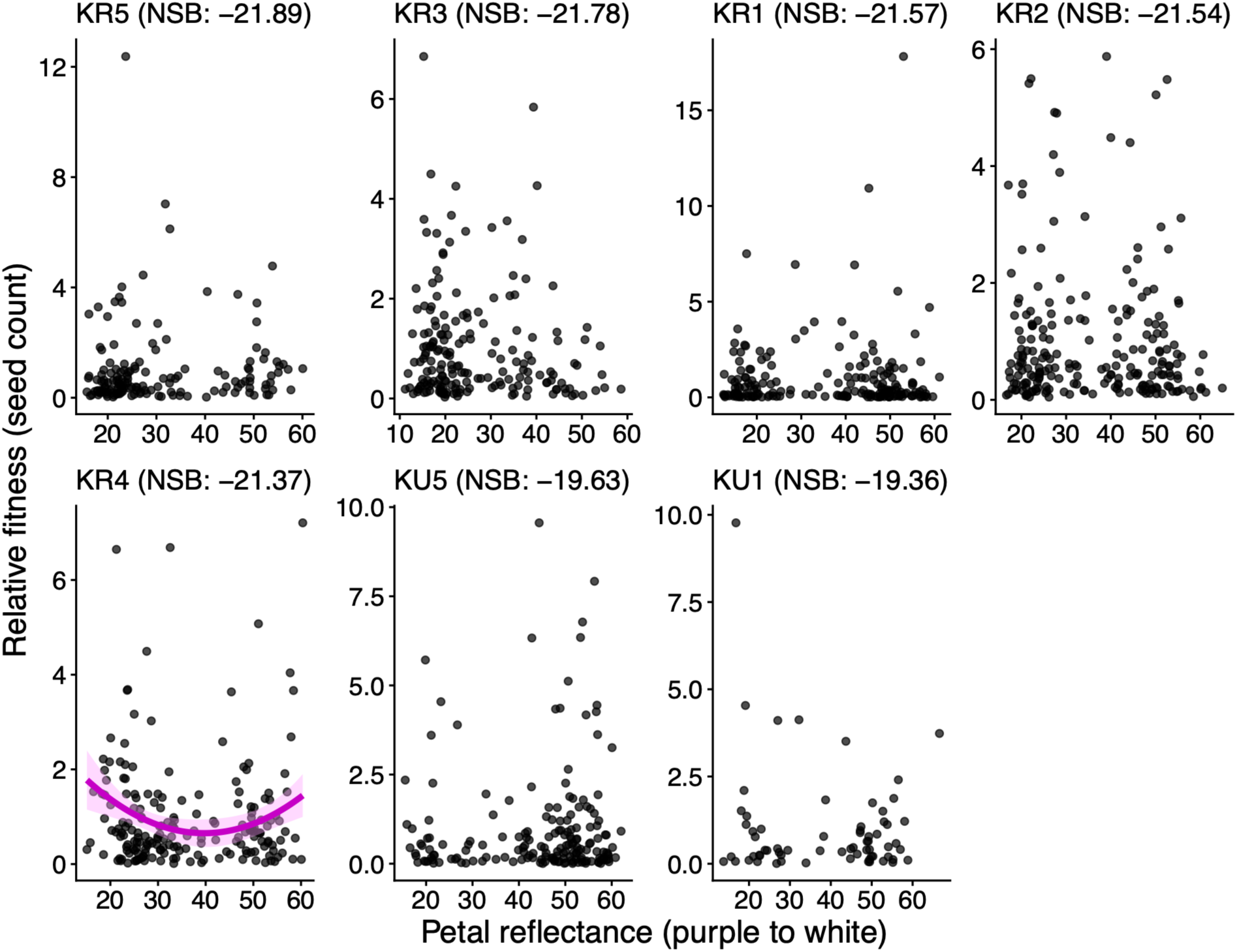
The relationship between relative fitness (the total seed count for each plant standardized by the population average) and petal reflectance for each population. The purple line indicates a significant quadratic regression. No linear regressions are shown because none were significant. Each point represents one plant. Petal reflectance is the average percentage of light reflected between 430 and 630 nm. Night sky brightness (NSB) is an estimate of human activity levels, where a more positive value represents a greater amount of human activity.

**Table 1:** Linear and quadratic selection differentials for the strength of selection on flower petal colour for each population. The linear selection differential (s) is equivalent to the slope coefficient of the linear regression between relative fitness and flower colour. The non-linear selection differential (C) is calculated by multiplying the coefficient of the squared term in the quadratic regression by two and represents the strength of disruptive or stabilizing selection acting on flower colour (Lande and Arnold 1983; Stinchcombe *et al*. 2008). The standard error (SE) for the quadratic selection differentials is also obtained by multiplying the SE of the squared term of the quadratic regression by two.

|  | Linear selection differential (s) |  |  | Quadratic selection differential (C) |  |  |
|---|---|---|---|---|---|---|
| Population | s | SE | p | C | SE | p |
| KR5 | 0.084228 | 0.119524 | 0.482057 | -0.342765 | 0.327785 | 0.297339 |
| KR3 | -0.100391 | 0.080801 | 0.215624 | -0.233449 | 0.171492 | 0.175072 |
| KR1 | 0.054113 | 0.138117 | 0.695653 | -0.425622 | 0.435102 | 0.329219 |
| KR2 | -0.060433 | 0.080524 | 0.453778 | -0.359221 | 0.221462 | 0.106266 |
| KR4 | 0.016857 | 0.085293 | 0.843541 | **0.658640** | **0.223160** | **0.003568** |
| KU5 | 0.066845 | 0.117514 | 0.570198 | 0.059532 | 0.279631 | 0.831656 |
| KU1 | -0.193572 | 0.186808 | 0.303945 | 0.956588 | 0.514716 | 0.067701 |

## Discussion

We investigated the influence of selection on a previously identified urban-rural cline in the flower colour of *H. matronalis* populations, in eastern Ontario (Maunder *et al*. 2026). We measured pollen limitation via a pollen supplementation experiment to investigate the potential for pollinators to impose selection on flower petal colour. We found almost no evidence that flowers are pollen limited and thus did not find evidence to support a possible role for pollinators as selective agents on flower colour. To assess if selection is meaningfully affecting the previously observed cline, we measured the strength of linear and non-linear selection on flower colour in populations across the urban-rural gradient. We found almost no evidence that flower colour is under selection and no evidence that selection is contributing to the observed urban-rural cline. Below we discuss the implications of our results from the pollen supplementation experiment and the strength of selection analysis.

Pollen limitation can be associated with stronger pollinator-mediated selection on floral traits (Ashman and Morgan 2004; Trunschke *et al*. 2017). Thus, if *H. matronalis* populations were pollen limited, it could indicate an increased opportunity in these populations for selection by pollinators. However, we did not find evidence that pollen supplementation improved the fitness of *H. matronalis* fruits and therefore did not find support for pollen limitation in these populations. However, since our pollen supplementation experiment only assessed pollen limitation at the level of pollen quantity, our results cannot speak to any possible limitations of pollen quality (Aizen and Harder 2007). Our pollen supplementation experiment was performed on a limited number of flowers and not the entire plant. When only a subset of flowers are given supplemental pollen, pollen limitation is often overestimated because resources may be reallocated between flowers to boost seed production in the fruits of supplemented flowers, while the plant’s overall seed production remains unchanged (Knight *et al*. 2006). Thus, if we found evidence of pollen limitation we would have to treat this finding with caution, but since we did not, this concern does not apply. Majetic *et al.’s* observation that pollinators did not show a colour preference in populations of *H. matronalis* in Pennsylvania (2008), aligns with our finding of limited potential for pollinator mediated selection. If *H. matronalis* is indeed not strongly pollen limited, there may be little opportunity for pollinator-mediated selection on any floral traits, including flower colour (Vanhoenacker *et al*. 2013; Trunschke *et al*. 2017).

After measuring the strength of selection acting on flower colour within populations, we found no evidence that selection is contributing to the observed cline. It is perhaps unsurprising that we did not find evidence of directional selection, since flower colour polymorphism in *H. matronalis* populations seem to be fairly well maintained over time (though temporal variation in directional selection can help maintain polymorphisms (Schmidt and Conde 2006; Fitzpatrick *et al*. 2007)). The earliest published report of naturalized *H. matronalis* populations in Canada mentions the polymorphism (Leon 1862) and observations of *H. matronalis* populations in London, ON found that the flower colour polymorphism was maintained in most populations over the span of forty years (Cavers unpublished – as mentioned in Francis *et al*. 2009). It is perhaps more surprising that we only found evidence of non-linear selection in one population, as both balancing and disruptive selection can contribute to the maintenance of phenotypic polymorphisms (Mitchell-Olds *et al*. 2007; McKinnon and Pierotti 2010; Kellenberger *et al*. 2019; Ito and Konuma 2020). More specifically, opposing selection between pollinators and other drivers (herbivores, pathogens, abiotic conditions, etc.) has been suggested as a mechanism that can maintain flower colour polymorphisms in other species (Irwin *et al*. 2003; Frey 2004; Fetterly *et al*. 2025).

While we did not detect selection on flower colour in these populations, this does not rule out that selection or human activity may be producing the urban rural cline. It is possible that because we focused on populations with a high diversity of flower colour (to facilitate adequate representation of white, pink, and purple individuals) that we inadvertently sampled populations where selection is weaker. Additionally, our estimate of fitness only reflects variation in potential seed set within one sampling year and does not reflect variation in other aspects of fitness such as survival prior to flowering or fitness due to pollen success. There are limited examples in the literature of other studies that found no evidence of selection acting on flower colour variation, though this is potentially influenced by publication bias against negative results (Rosenthal 1979). For example, Keasar *et al*. did not find evidence to support an adaptive interpretation for clinal variation in the flower colour frequency of *Silybum marianum* (2016) and Parachnowitsch and Kessler found no significant directional selection on flower colour in *Penstemon digitalis* (2010). In the context of pollinator-mediated selection, a meta-analysis found that even when selection on flower colour is significant, it is generally weaker than selection on floral traits that affect pollinator efficiency, rather than pollinator attraction (Caruso *et al*. 2018). Altogether, our results highlight the possibility that selection may not be an important contributor to the previously observed urban-rural cline in flower colour.

## Conclusions and Future Directions

We encourage future research to investigate the influence that non-selective and historical processes may have on the observed cline. Given the historical context of *H. matronalis*’s introduction as a garden plant, multiple non-adaptive processes could contribute to the observed cline: i) historical bias in flower colour during introduction, ii) multiple introductions increasing variation in urban areas, or iii) genetic drift during expansion (see Maunder *et al*. 2026). To assert a role for neutral processes on the observed cline, an investigation into neutral genetic variation is required (Wang *et al*. 2016). Recent research on *H. matronalis* populations across two transects in the Great Lakes region found almost no neutral genetic structure between populations (Johnson *et al*. 2025). If this finding extends to the populations in our study region, it makes the contribution of non-adaptive processes uncertain. It is possible that while no neutral structure exists between populations at a large scale (between cities), that there may still be neutral population genetic structure at a smaller scale (increased distance from one city).

## Data Accessibility Statement

Data and analyses are openly available in Zenodo at https://doi.org/10.5281/zenodo.24040006

## Author Contributions

KGM, JRS, and MB conceived the project. KGM and MGM conducted fieldwork. KGM analyzed the data. JRS and MB provided supervision and training. KGM wrote the paper with help from all co-authors.

## Acknowledgements

We thank Louisa Bartkovich and Martina Gjevori for their assistance in the field and Claire Abbasi, Cynthi Ni, Ellie Hector, Emma Zongaro, Francesca Tuazon, Jo Fletcher, Jon Agolli, Julia Cruz, Katie Monat, Lilyan Alassaf, Maura McGregor, Nora Hand-Lennon, Osmond Hui, Shayne Enriquez, Stephanie Leung, and Tara McGruder for the hours spent counting and measuring fruit. This work was supported by NSERC Discovery Grants to JRS and MB; awards from the Canada Foundation for Innovation John R. Evans Leaders Fund and the Ontario Research Fund to MB; NSERC CGS-D, OGS, and UofT FAST doctoral fellowships to KGM; and a CGCS summer fellowship to MGM.

## Appendix A Lower proportion of intact seeds in urban populations

All marked fruits from the supplemental pollen experiment were opened and the number of intact seeds within fruits was counted. We defined “intact seeds” as those that could withstand being rolled between two fingers with gentle force. The majority of potential seeds, if still in one piece after the fruit was split open, crumbled very easily and did not meet this standard. We calculated the proportion of seeds that remained intact by dividing the number of intact seeds within a fruit by our estimate of total potential seed number (fruit length multiplied by 2.06, see Appendix B Figure 1). In a few cases this calculation produced a value greater than 1 and so we truncated this measure and assigned those individuals a value of 1.

We investigated whether the proportion of intact seeds within fruits was impacted by human activity, reflectance, or the pollen treatment. To do this, we performed a zero-one inflated beta (ZOIB) mixed regression using the brms package (Bürkner 2017). The script for the ZOIB regression model was initially generated using DeepSeek-V3.2 (accessed April 2026) and was then reviewed and edited by the authors to confirm accuracy and suitability. For the ZOIB model, the proportion of intact seeds within a fruit was used as the dependent variable; human activity, reflectance (z-score normalised), and pollen treatment as fixed effects; and plant identity as a random effect. We analyzed how these factors affected the predicted mean proportion of intact seeds (for continuous values between 0 and 1), as well as the zero-or-one inflation (zoi) and the conditional one inflation (coi).

The mean proportion of intact seeds within fruits was fairly low at just 27%. The mean proportion of intact seeds was negatively associated with human activity (Appendix A Figure 1; posterior mean estimate: -0.17, CrI: [-0.25, -0.10), corresponding to a change from 33% intact seeds at the lowest level of human activity (-21.89 mag/arc sec^2^) and 23% intact seeds at the highest human activity level (-19.17 mag/arc sec^2^). The 95% credible intervals for the association between the mean proportion of intact seeds and both reflectance and pollen supplementation overlapped with zero, as did the association of all variables on zoi and coi (Appendix A Table 1).

We are not alone in observing a relatively low rate of intact seeds within *H. matronalis* fruits. Previous research in our study area similarly noted high levels of seed destruction and found evidence that the seed weevil *Ceutorhynchus inaffectatu*s was responsible (Ensing *et al*. 2025). This weevil was first recorded in North America in 2018 (five years before this study).

However, the low number of intact seeds could also represent early ovule abortion which has been previously observed at high frequency in this species (Susko and Clubb 2008). While our estimates of plant fitness do not include a consideration of whether seeds were intact, we did not find any association between flower colour and the proportion of intact seeds (Appendix A Table 1), so this should not affect our selection estimates. We did however identify a significant decrease in the proportion of intact seeds in populations in more urban areas. If a lack of intact seeds is due to seed predation this could indicate a higher presence or higher destruction level of the *C. inaffectatu*s in urban areas. If a lack of intact seeds is due to early ovule abortion, our results could reflect a decrease in resource quality or availability in more urban populations, as Susko and Club suspected that resource competition was likely responsible for early ovule abortion (2008).

**Appendix A Figure 1:**
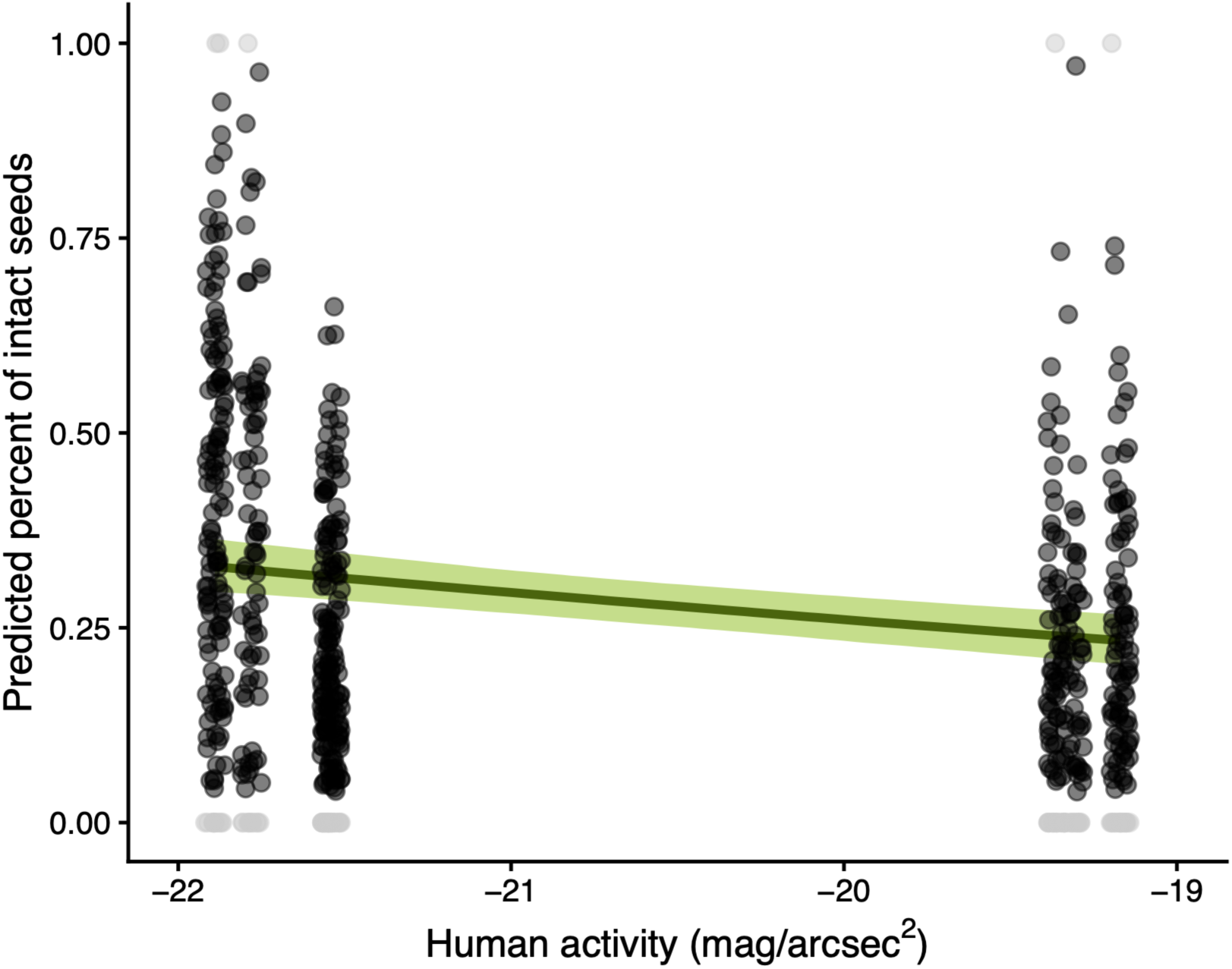
A decrease in the proportion of intact seeds within fruits as the amount of human activity around a population increases. Each point represents an individual fruit, with 1-3 fruits per plant. The trend line and ribbon are from predictions from zero-one-inflate beta regression. The light grey points indicate fruits where either none or all seeds were intact; these fruits are not included in the shown beta regression (but were included in calculations of zoi and/or coi).

**Appendix A Table 1:** Summary of the zero-one inflated beta (ZOIB) mixed regression model. Formulas: Proportion of intact seeds ∼ NSB + scale(reflectance) + pollen supplementation + (1|plant identity); zoi ∼ NSB + scale(reflectance) + pollen supplementation + (1|plant identity); coi ∼ NSB + scale(reflectance) + pollen supplementation + (1|plant identity). We used weakly informative priors, specifically we used student_t(3, 0, 2.5) for the random effect and normal(0, 1) for mean, zoi, and coi. We estimated the model with four chains, 2000 iterations per chain, and 4000 post-warmup draws.

| Fixed Effects |  |  |  |  |  |  |
|---|---|---|---|---|---|---|
| Sub-models | Parameter | Estimate | Est.Error | Lower 95% CrI | Upper 95% CrI | Bulk_ESS |
| Mean | Intercept | -4.52 | 0.79 | -6.06 | -2.98 | 2428 |
|  | **NSB** | **-0.17** | **0.04** | **-0.25** | **-0.1** | **2393** |
|  | scale(reflectance) | 0 | 0.04 | -0.09 | 0.08 | 2511 |
|  | pollen supplementation | 0 | 0.09 | -0.18 | 0.18 | 2470 |
| Zoi | Intercept | 2.17 | 2.29 | -2.36 | 6.74 | 3398 |
|  | NSB | 0.21 | 0.11 | 0 | 0.44 | 3166 |
|  | scale(reflectance) | 0.08 | 0.13 | -0.17 | 0.34 | 3445 |
|  | pollen supplementation | 0.42 | 0.26 | -0.08 | 0.95 | 3526 |
| Coi | Intercept | -8.56 | 8.99 | -28.4 | 7.34 | 4095 |
|  | NSB | -0.24 | 0.43 | -1.15 | 0.52 | 4274 |
|  | scale(reflectance) | -0.02 | 0.47 | -0.97 | 0.92 | 4294 |
|  | pollen supplementation | -0.14 | 0.73 | -1.56 | 1.38 | 4313 |
| Random Effect |  |  |  |  |  |  |
|  |  | Estimate | Est.Error | Lower 95% CrI | Upper 95% CrI | Bulk_ESS |
| Mean | SD (intercept) | 0.51 | 0.06 | 0.4 | 0.62 | 1048 |
| Zoi | SD (intercept) | 1 | 0.31 | 0.31 | 1.58 | 594 |
| Coi | SD (intercept) | 1.21 | 0.97 | 0.05 | 3.57 | 762 |

## Appendix B Additional figures

**Appendix B Figure 1:**
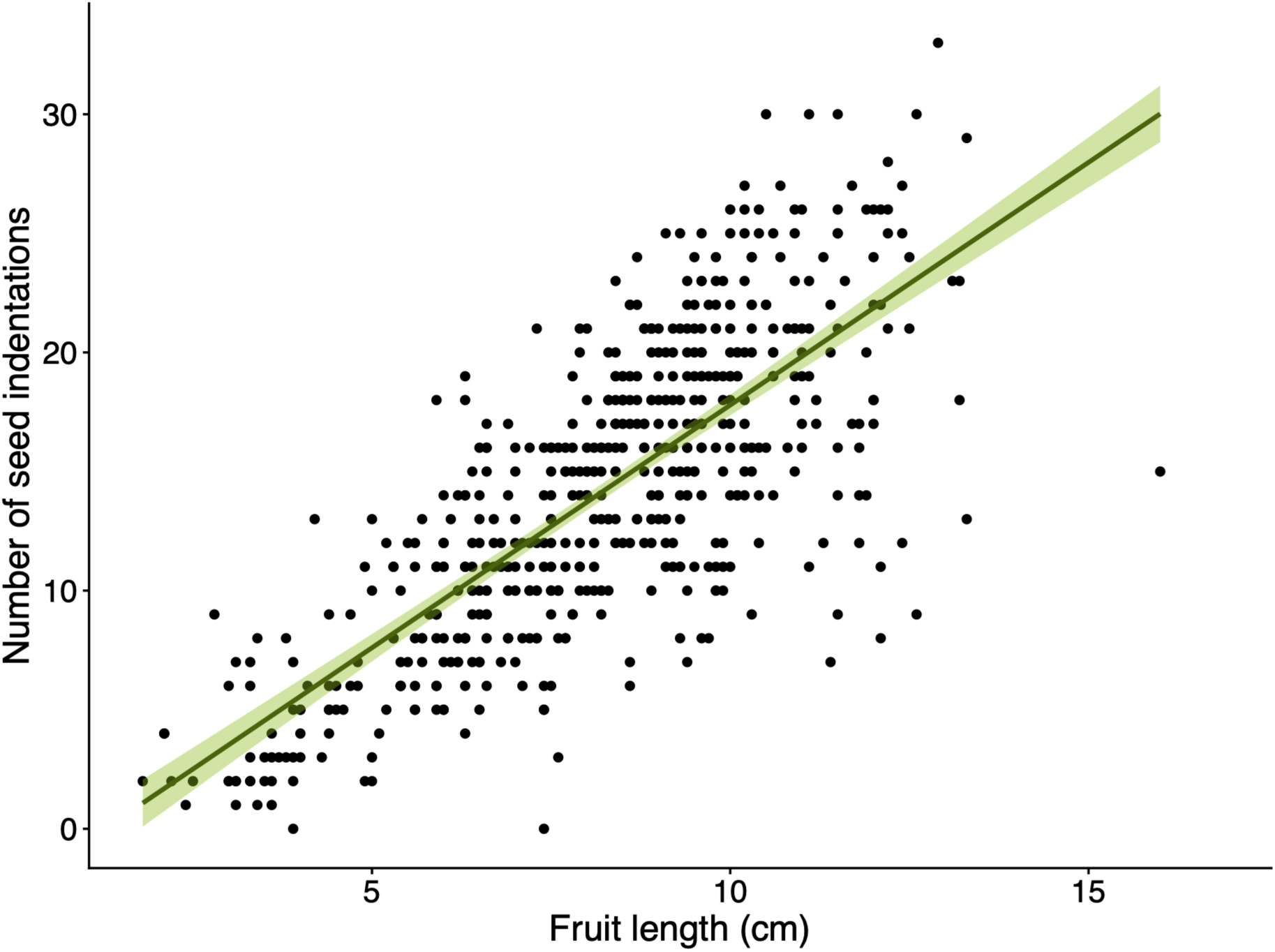
Linear relationship between fruit length and the number of seed indentations within a fruit. To obtain this data, all marked fruits from the supplemental pollen experiment were opened, and the number of seed indentations along the septum was counted (evidence of seed development (Mitchell and Ankeny 2001)). We ran a linear mixed effect model with the number of seed indentations as the dependent variable, fruit length as a fixed effect, and plant identity as a random effect (to account for the fact that as many as three fruits may be from the same plant). We found a significant positive association between number of seed indentations and fruit length (slope = 2.059, df = 563.6, p < 0.001). Given this fairly tight relationship, for our fitness analysis we estimated the potential number of seeds within a fruit by multiplying fruit length by 2.06.

**Appendix B Figure 2:**
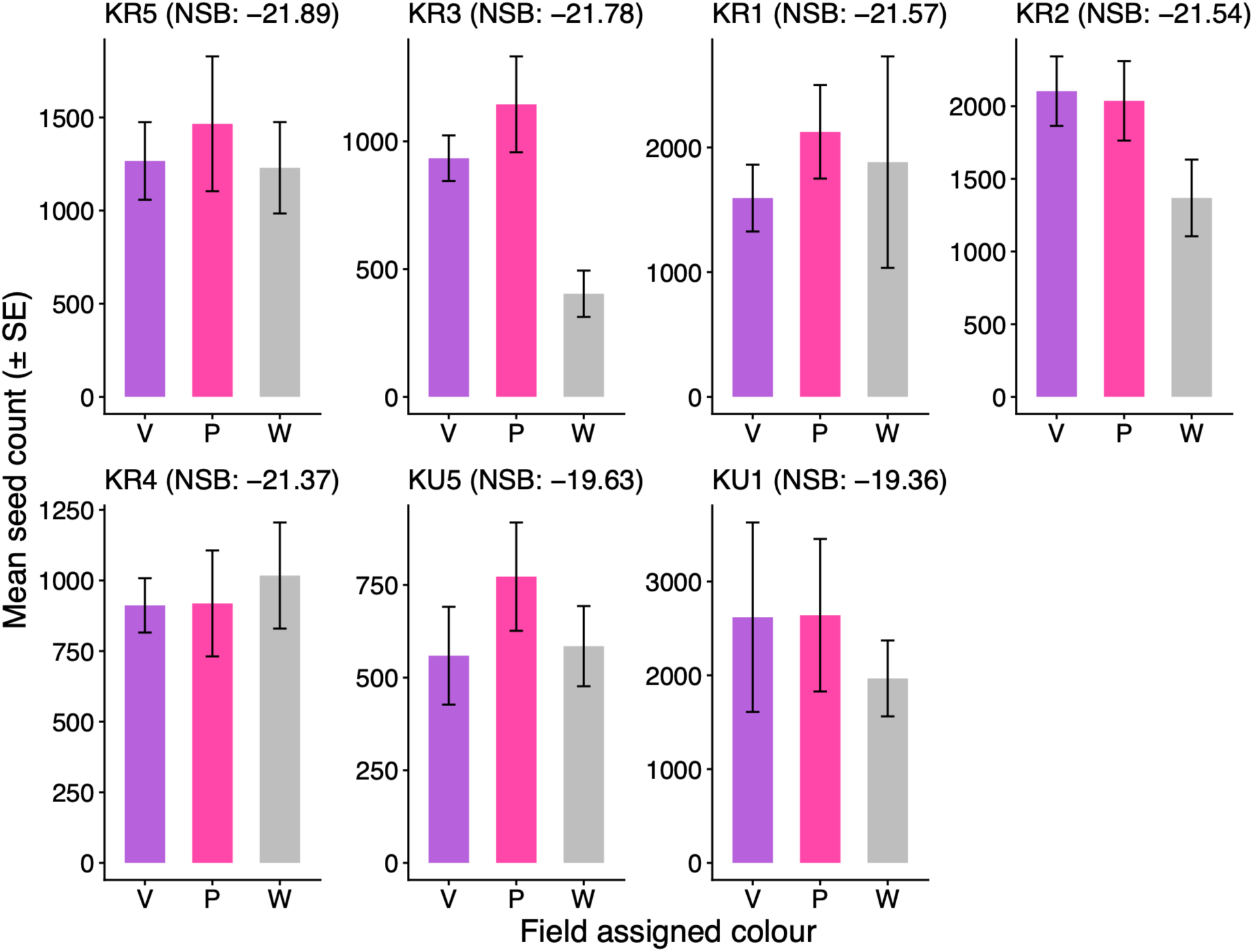
Variation in absolute fitness (potential seed count for the whole plant) between populations and field assigned colour groups (V = purple, P = pink, W = white). We ran a generalized linear model with a gamma distribution with absolute fitness as the dependent variable and for independent variables used population, colour, and their interaction. We ran a type III test and found that there was a significant main effect of population (χ² = 32.01, df = 6, p = < 0.001), but not a significant effect of colour (p = 0.46) or the interaction of population and colour (0.71). The black bars represent standard error.

## Literature Cited

Aizen MA, Harder LD. 2007. Expanding the limits of the pollen-limitation concept: Effects of pollen quantity and quality. Ecology 88: 271–281.

Al-Ghussain L. 2019. Global warming: review on driving forces and mitigation. Environmental Progress and Sustainable Energy 38: 13–21.

Arista M, Talavera M, Berjano R, Ortiz PL. 2013. Abiotic factors may explain the geographical distribution of flower colour morphs and the maintenance of colour polymorphism in the scarlet pimpernel. Journal of Ecology 101: 1613–1622.

Ashman TL, Morgan MT. 2004. Explaining phenotypic selection on plant attractive characters: male function, gender balance or ecological context? Proceedings of the Royal Society B: Biological Sciences 271: 553.

Bode RF, Tong R. 2018. Pollinators exert positive selection on flower size on urban, but not on rural Scotch broom (*Cytisus scoparius L*.). Journal of Plant Ecology 11: 493–501.

Buchholz S, Kowarik I. 2019. Urbanisation modulates plant-pollinator interactions in invasive vs. native plant species. Scientific Reports 9: 6375.

Bürkner, P.-C. 2017. Brms: An R package for Bayesian multilevel models using Stan. Journal of Statistical Software, 80: 1–28. 10.18637/jss.v080.i01.

Capua G Di, Rahmstorf S. 2023. Extreme weather in a changing climate. Environmental Research Letters 18: 102001.

Caruso CM, Eisen KE, Martin RA, Sletvold N. 2018. A meta-analysis of the agents of selection on floral traits. Evolution 73: 4–14.

Charmantier A, Burkhard T, Gervais L, Perrier C, Schulte-Hostedde AI, Thompson MJ. 2024. How does urbanization affect natural selection? Functional Ecology 38: 2522–2536.

Ensing DJ, Nelson TD, Moffat CE, et al. 2025. Together again: the invasive mustard *Hesperis matronalis* suffers devastating seed predation by a recently adventive specialist weevil. BioControl 70: 835–847.

Falchi F, Cinzano P, Duriscoe D, et al. 2016. The new world atlas of artificial night sky brightness. Science Advances 2: e1600377.

Fetterly E, Braum AS, Kim C, et al. 2025. Selection maintains floral color polymorphism in scarlet paintbrush, Castilleja coccinea, reflecting combined ecological factors. American Journal of Botany: e70094.

Fishman L, Willis JH. 2008. Pollen limitation and natural selection on floral characters in the yellow monkeyflower, *Mimulus guttatus*. New Phytologist 177: 802–810.

Fitzpatrick MJ, Feder E, Rowe L, Sokolowski MB. 2007. Maintaining a behaviour polymorphism by frequency-dependent selection on a single gene. Nature 447: 210–212.

Francis A, Cavers PB, Warwick SI. 2009. The biology of Canadian weeds. 140. Hesperis matronalis L. Canadian Journal of Plant Science 89: 191–206.

Frey FM. 2004. Opposing natural selection from herbivores and pathogens may maintain floral-color variation in *Claytonia virginica* (Portulacaceae). Evolution 58: 2426–2437.

Glover BJ, Martin C. 1998. The role of petal cell shape and pigmentation in pollination success in *Antirrhinum majus*. Heredity 1998 80:6 80: 778–784.

Grimm NB, Faeth SH, Golubiewski NE, et al. 2008. Global change and the ecology of cities. Science 319: 756–760.

Hao L, Sun G, Huang X, et al. 2023. Urbanization alters atmospheric dryness through land evapotranspiration. npj Climate and Atmospheric Science 6: 149.

Harrison T, Winfree R. 2015. Urban drivers of plant-pollinator interactions. Functional Ecology 29: 879–888.

Irwin RE, Strauss SY, Storz S, Emerson A, Guibert G. 2003. The role of herbivores in the maintenance of flower colour polymorphism in wild radish. Ecology 84: 1733–1743.

Irwin RE, Warren PS, Adler LS. 2018. Phenotypic selection on floral traits in an urban landscape. Proceedings of the Royal Society B: Biological Sciences 285: 20181239.

Ito S, Konuma J. 2020. Disruptive selection of shell colour in land snails: a mark–recapture study of *Euhadra peliomphala simodae*. Biological Journal of the Linnean Society 129: 323–333.

De Jager ML, Ellis AG. 2014. Floral polymorphism and the fitness implications of attracting pollinating and florivorous insects. Annals of Botany 113: 213–222.

Johnson SE, Rifkin JL, Wright SI, Baucom RS. 2025. Color, size, shape: The drivers of floral variation in *Hesperis matronalis* (Dame’s Rocket). bioRxiv: 2025.10.16.682931.

Kalnay E, Cai M. 2003. Impact of urbanization and land-use change on climate. Nature 423: 528–531.

Keasar T, Gerchman Y, Lev-Yadun S. 2016. A seven-year study of flower-color polymorphism in a Mediterranean annual plant. Basic and Applied Ecology 17: 741–750.

Kellenberger RT, Byers KJRP, De Brito Francisco RM, et al. 2019. Emergence of a floral colour polymorphism by pollinator-mediated overdominance. Nature Communications 10: 63.

Keller SR, Taylor DR. 2008. History, chance and adaptation during biological invasion: separating stochastic phenotypic evolution from response to selection. Ecol. Lett. 11: 852–866.

Knight TM, Steets JA, Ashman TL. 2006. A quantitative synthesis of pollen supplementation experiments highlights the contribution of resource reallocation to estimates of pollen limitation. American Journal of Botany 93: 271–277.

Kuznetsova, A., Brockhoff, P. B., & Christensen, R. H. B. 2017. lmerTest package: tests in linear mixed effects models. Journal of Statistical Software, 82: 1–26. 10.18637/jss.v082.i13.

Lande R, Arnold SJ. 1983. The measurement of selection on correlated characters. Evolution 37: 1210–1226.

Leon P. 1862. Le Flore canadienne; ou, Description de toutes les plantes des forêts, champs, jardins et eaux du Canada, donnant le nom botanique de chacune, ses noms vulgaires français et anglais, indiquant son parcours géographique, les propriétés qui la distinguent. Quebec: Joseph Darveau.

Li T, Shilling F, Thorne J, et al. 2010. Fragmentation of China’s landscape by roads and urban areas. Landscape Ecology 25: 839–853.

Maia, R., Gruson, H., Endler, J. A., & White, T. E. 2019. pavo 2: New tools for the spectral and spatial analysis of colour in r. Methods in Ecology and Evolution, 10: 1097–1107. 10.1111/2041-210X.13174.

Majetic CJ. 2008. The evolutionary ecology of floral scent in Hesperis matronalis: assessing the potential for pollinator-mediated natural selection.

Maunder KG, Dawson K, Joslin L, et al. 2026. Random and non-random variation in flower colour along an urban–rural gradient in the introduced mustard *Hesperis matronalis*. Annals of Botany 137: 1290–1304.

McKinnon JS, Pierotti MER. 2010. Colour polymorphism and correlated characters: Genetic mechanisms and evolution. Molecular Ecology 19: 5101–5125.

McLean P, Gallien L, Wilson JRU, Gaertner M, Richardson DM. 2017. Small urban centres as launching sites for plant invasions in natural areas: insights from South Africa. Biological Invasions 19: 3541–3555.

Miles LS, Breitbart ST, Wagner HH, Johnson MTJ. 2019. Urbanization shapes the ecology and evolution of plant-arthropod herbivore interactions. Frontiers in Ecology and Evolution 7: 469218.

Mitchell RJ, Ankeny DP. 2001. Effects of local conspecific density on reproductive success in Penstemon digitalis and *Hesperis matronalis*. Ohio J. Sci. 101: 22–27.

Mitchell-Olds T, Willis JH, Goldstein DB. 2007. Which evolutionary processes influence natural genetic variation for phenotypic traits? Nature Reviews Genetics: 845–856.

Parachnowitsch AL, Kessler A. 2010. Pollinators exert natural selection on flower size and floral display in *Penstemon digitalis*. New Phytologist 188: 393–402.

Phelan PE, Kaloush K, Miner M, et al. 2015. Urban heat island: mechanisms, implications, and possible remedies. Annual Review of Environment and Resources 40: 285–307.

Des Roches S, Brans KI, Lambert MR, et al. 2021. Socio-eco-evolutionary dynamics in cities. Evolutionary Applications 14: 248–267.

R Core Team 2024. R: a language and environment for statistical computing. R Foundation for Statistical Computing, Vienna, Austria. https://www.R-project.org/.

Rosenthal R. 1979. The file drawer problem and tolerance for null results. Psychological Bulletin 86: 638–641.

Sandring S, Ågren J. 2009. Pollinator-mediated selection on floral display and flowering time in the perennial herb *Arabidopsis lyrata*. Evolution 63: 1292 – 1300.

Schemske DW, Bierzychudek P. 2001. Perspective: Evolution of flower color in the desert annual *Linanthus parryae*: Wright revisited. Evolution 55: 1269–1282.

Schmidt PS, Conde DR. 2006. Environmental heterogeneity and the maintenance of genetic variation for reproductive diapause in drosophila melanogaster. Evolution 60: 1602–1611.

Stinchcombe JR, Agrawal AF, Hohenlohe PA, Arnold SJ, Blows MW. 2008. Estimating nonlinear selection gradients using quadratic regression coefficients: double or nothing? Evolution 62: 2435–2440.

Strauss S, Whittall J. 2007. Non-pollinator agents of selection on floral traits In: Lawrence D. Harder, Spencer C. H. Barrett, eds. Ecology and Evolution of Flowers. Oxford University Press, 120–138.

Susko DJ, Clubb M. 2008. Pollination effects on patterns of ovule fate in *Hesperis matronalis* (Brassicaceae). Botany 86: 466–474.

Tenhumberg B, Dellinger AS, Smith SD. 2023. Modelling pollinator and nonpollinator selection on flower colour variation. Journal of Ecology 111: 746–760.

Theodorou P, Radzevičiūtė R, Lentendu G, et al. 2020. Urban areas as hotspots for bees and pollination but not a panacea for all insects. Nature Communications 11: 576.

Trunschke J, Sletvold N, Ågren J. 2017. Interaction intensity and pollinator-mediated selection. New Phytologist 214: 1381–1389.

Vanhoenacker D, Ågren J, Ehrlén J. 2013. Non-linear relationship between intensity of plant-animal interactions and selection strength. Ecology Letters 16: 198–205.

Wang H, Talavera M, Min Y, Flaven E, Imbert E. 2016. Neutral processes contribute to patterns of spatial variation for flower colour in the Mediterranean *Iris lutescens* (Iridaceae). Annals of Botany 117: 995–1007.

Weeks EL, Frey FM. 2007. Seed production and insect visitation rates in *Hesperis matronalis* are not affected by floral symmetry. International Journal of Plant Sciences 168: 611–617.

Wenzel A, Grass I, Belavadi V V., Tscharntke T. 2020. How urbanization is driving pollinator diversity and pollination – A systematic review. Biological Conservation 241: 108321.

